# Multiplex-inhibitor bead mass spectrometry reveals differential kinome signatures in newly diagnosed and recurrent glioblastoma

**DOI:** 10.1101/2020.09.21.306910

**Authors:** Anna Jermakowicz, Alison M. Kurimchak, Jann Sarkaria, Ricardo Komotar, Michael E. Ivan, Stephan Schürer, James S. Duncan, Nagi G. Ayad

## Abstract

Glioblastoma (GBM) is the most common and aggressive adult brain tumor. Despite years of research, clinical trials have not improved the outcome for GBM. Standard of care for newly diagnosed GBM includes surgical resection, followed by radiation and chemotherapy. Tumor recurrence is inevitable and since most patients are not candidates for a second surgical resection, there is an urgent need to identify resistance mechanisms that arise in recurrent GBM. We postulated that examining the differences of activated kinases between newly diagnosed and recurrent GBM may provide insight to resistance mechanisms.

To map the kinome landscape of newly diagnosed (nGBM) and recurrent GBM (rGBM) patient derived xenograft tumors, we used Multiplexed Inhibitor Beads and Mass Spectrometry (MIB-MS). We performed pathway analysis of kinases that differed in MIB-binding between nGBM and rGBM to identify kinase-driven signaling pathways. We also analyzed transcriptional profiles to determine the overlap in signaling pathways seen using proteomics or transcriptomics.

Using MIB-MS kinome profiling, we found key differences in kinase-driven signaling pathways that may account for the increase in aggressive behavior seen in recurrent GBM. This included a shift in pathways driving cell invasion and proliferation, as well as upregulation of signaling pathways that drive GBM stem-cell like cell differentiation. Analysis of RNA-sequencing showed no statistically significant differences between enriched gene ontologies in nGBM and rGBM, demonstrating the importance of MIB-MS kinome profiling. Collectively, these studies suggest that kinome profiling may inform future clinical trials for kinase inhibitors in GBM.

## INTRODUCTION

Glioblastoma (GBM) accounts for 50 percent of patients with gliomas, making it the deadliest form of brain cancer ^1^. The standard of care for primary treatment is surgical resection followed by radiation and temozolomide (TMZ) chemotherapy, with a median progression free survival of 6.9 months and median overall survival of 14.6 months ^2^. However, complete resection is impossible due to the diffuse infiltration of GBM cells into the surrounding brain tissue and nearly universal resistance to both radiation and TMZ ^3,4^. Unfortunately, little progress has been made to advance therapies, and the current 5-year survival is 9.8% ^2^.

The treatment of recurrent GBM is particularly difficult. Tumor growth is invasive and the use of many conventional chemotherapeutic agents is limited due to their toxicity in the brain and inability to cross the blood-brain barrier ^1^. Tumor recurrence typically occurs within 2 cm of the resection cavity, yet only one in four patients is a candidate for repeat resection ^3,5,6^. Although preliminary studies suggest that combination therapy with targeted agents may increase tumor radiosensitivity, ongoing clinical trials have shown minimal survival benefit ^6-8^. Additionally, radiotherapy may paradoxically promote tumor growth in these patients, particularly when given in doses insufficient to irradiate tumor cells ^4,9^. As a result, there is currently no standard of care for recurrent GBM where patients either receive palliative care or are enrolled in clinical trials ^4^.

Radiation and chemotherapy are important adjuvant therapies for GBM, however resistance to both is universal. Radiation induces double stranded breaks in DNA, with the goal of overwhelming the DNA mismatch repair system, leading to apoptosis ^4^. However, radiation therapy cannot be safely administered at doses sufficient to irradiate the entire tumor cell population, contributing to frequent recurrence near the tumor resection cavity ^3^. The irradiated tumor microenvironment is more highly vascularized and necrotic, making it an ideal environment for tumor recurrence ^10^. Additionally, recurrent GBM tumors are more aggressive as a result of radiation therapy-induced HIF-1 activation and upregulation of VEGF, which leads to increased immunosuppression, decreased apoptosis of tumor cells, and increased cell motility ^4,5,11^. TMZ is a chemotherapeutic agent that alkylates and methylates DNA, leading to DNA damage and death of tumor cells. However, GBM rapidly gains resistance to TMZ due to either an inherent or acquired overexpression of the O6-metylguanine DNA methyltransferase (MGMT) DNA repair, overexpression of epidermal growth factor receptor (EGFR), or restoration of p53 activity by Mdm2 inhibition ^12,13^.

Overexpression of receptor tyrosine kinases (RTKs) is prevalent among many cancers, including GBM, and leads to enhanced cellular migration, proliferation, angiogenesis, and tumorigenesis ^14^. EGFR is one such RTK that is overexpressed in 40% of GBM tumors. Following TGFα and EGF ligand binding, EGFR is activated by dimerization ^13,15^. Activated EGFR drives downstream PI3K/AKT/mTOR, RAS/RAF/MAPK, and JAK/STAT pathways. Of particular importance is the PI3K/AKT/mTOR pathway, which leads to an elevation in c-MYC, resulting in the promotion of tumorigenesis through anaerobic glycolysis ^13^.

Given the heterogeneity among both GBM tumors and GBM cell populations, as well as their ability to rapidly acquire drug resistance, a one-size-fits-all treatment approach for recurrent GBM is not possible ^16^. Unfortunately, differences in newly diagnosed and recurrent GBM tumors are difficult to differentiate based on mRNA levels alone, suggesting a need for further exploration of tumor signaling pathways. Since the majority of GBM tumors have alterations in RTK signaling, we believe that kinase expression and activity may provide insight into the changes in tumorigenesis pathways. Previously, phosphoproteomics has been used to characterize signaling in GBM, providing important knowledge of specific phosphosites driving kinase activation and the effects of kinase inhibitors ^17^. Kinome proteomic profiling has been used recently to determine kinases that are upregulated or downregulated in different cancer types and to assess the kinome reprogramming induced by kinase inhibitors ^18-23^. Since the activity of kinases in not well correlated to transcriptional data, we used Multiplexed Inhibitor Beads coupled with Mass Spectrometry (MIB-MS) to provide a snapshot of the tumor kinome profiles in a panel of newly diagnosed and recurrent GBM tumors. Protein kinases are enriched from lysates by immobilized kinase inhibitor-resins based on their affinity, abundance, and in some instances activity ^18^. Utilizing this method, we report differences in MIB-binding of several kinases in newly diagnosed GBM relative to recurrent GBM. We demonstrate that the MAP kinase pathway is preferentially upregulated in newly diagnosed GBM. These studies provide one of the first indications that there are kinome wide differences between GBM in the newly diagnosed and recurrent settings, which can inform future clinical trials utilizing kinase inhibitors for the treatment of GBM.

## METHODS

### Patient derived xenograft glioblastoma tumors

PDX Tumors were obtained from the Mayo Clinic Brain Tumor PDX national resource or from the University of Miami Brain Tumor Initiative ^24^. At the time of patient neurosurgical tumor resection, tissue is mixed with Matrigel and directly implanted into the flank of Crl:NU-*Foxn1*^*nu*^ mice. After tumor has reached 1 to 1.5 cm in size, the tumor is serial passaged to establish a tumor line. Established tumors were isolated from the flank of Crl:NU-*Foxn1*^*nu*^ mice and cryo-preserved in freezing media containing 10% DMSO prior to kinase extraction.

### GBM tumor active kinome proteomic profiling

We profiled the kinome in a panel of newly diagnosed and recurrent GBM PDX tumors at baseline using MIB-MS profiling, as previously described ^21^. For proteomic measurement of kinases in tissues, we used MIB-MS profiling and quantitated kinase levels using a super-SILAC (s-SILAC) ^25^. Briefly, an equal amount of s-SILAC reference (5 mg) was spiked into each primary tissue sample (5 mg) and kinases purified from tissues using MIB-resins, kinases were eluted, digested, and peptides analyzed by LC-MS/MS as previously described ^26^. To identify kinases differentially expressed in newly diagnosed or recurrent GBM tumors isolated from PDX models, we performed MIB-MS analysis on 6 newly diagnosed and 4 recurrent GBM tumors. Measurement of MIB-enriched kinase levels in tissues was performed s-SILAC quantitation using MaxQuant software version 1.6.1.0.

### Network analysis of MIB-MS kinome signatures

Differential MIB-binding of kinases was determined by an ANOVA (BH P≤0.05) and statistically significant kinases uploaded to STRING database (https://string-db.org/ version 11) for further analysis. Kinase protein interactions with medium confidence interactions based on experiments, databases, co-expression, neighborhood, gene fusion, and co-occurrence were mapped using k means clustering. KEGG pathway analysis was performed and significant pathways (FDR < 0.05) were identified.

### Differentially expressed genes and gene ontologies determined using RNA-sequencing

Normalized RNA-sequencing data from PDX GBM cell lines was downloaded form cBioPortal (https://www.cbioportal.org/study/summary?id=gbm_mayo_pdx_sarkaria_2019). Samples were normalized using log transformed RNA-seq RPKM z-scores relative to all samples. Differentially upregulated genes (z-score > 1.2) were identified for each cell line. Significantly upregulated genes for each cell line were uploaded to DAVID (https://david.ncifcrf.gov/) for biological process gene ontology enrichment analysis. Ontologies with a Benjamini corrected p-value < 0.05 were included for further analysis. Ontologies were compared between nGBM and rGBM using a two-way ANOVA (BH <0.05) of changes in fold enrichment for each pathway.

### Kinases predicted to be activated by mRNA levels

Differentially upregulated genes for GBM PDX tumors were uploaded to the eXpression2Kinases (X2K) platform (https://amp.pharm.mssm.edu/X2K/). This platform searches for transcription factors known to upregulate differentially expressed genes then computes a subnetwork of protein-protein interactions that connect those transcription factors, followed by a kinase enrichment analysis for the proteins involved in the subnetwork ^27,28^. Kinases predicted to be activated for each cell line were identified and statistically different kinases between nGBM and rGBM were calculated using two-way ANOVA (BH < 0.05). X2K kinases were uploaded to DAVID for KEGG pathway analysis. X2K kinases were visualized on a heatmap using -log(FDR) values. Kinases were also uploaded to STRING for network analysis and KEGG pathway enrichment analysis.

## RESULTS

### Kinome profiling reveals distinct kinase targets in newly diagnosed and recurrent GBM tumors

To explore the differences in kinome profiles of newly diagnosed (nGBM) and recurrent GBM (rGBM), we used MIB-MS. Six newly diagnosed and four recurrent GBM PDX tumors were isolated from the flank of Crl:NU-*Foxn1*^*nu*^ mice and kinases were extracted. Two-way ANOVA with Benjamini-Hochberg (BH) corrected p-values < 0.05 was used to identify kinases differentially activated in newly diagnosed or recurrent GBM. The analysis shows distinct kinase targets between newly diagnosed and recurrent tumors, providing us with a robust kinome signature in newly diagnosed versus recurrent GBM (Figure 1A-B).

**Figure 1.**
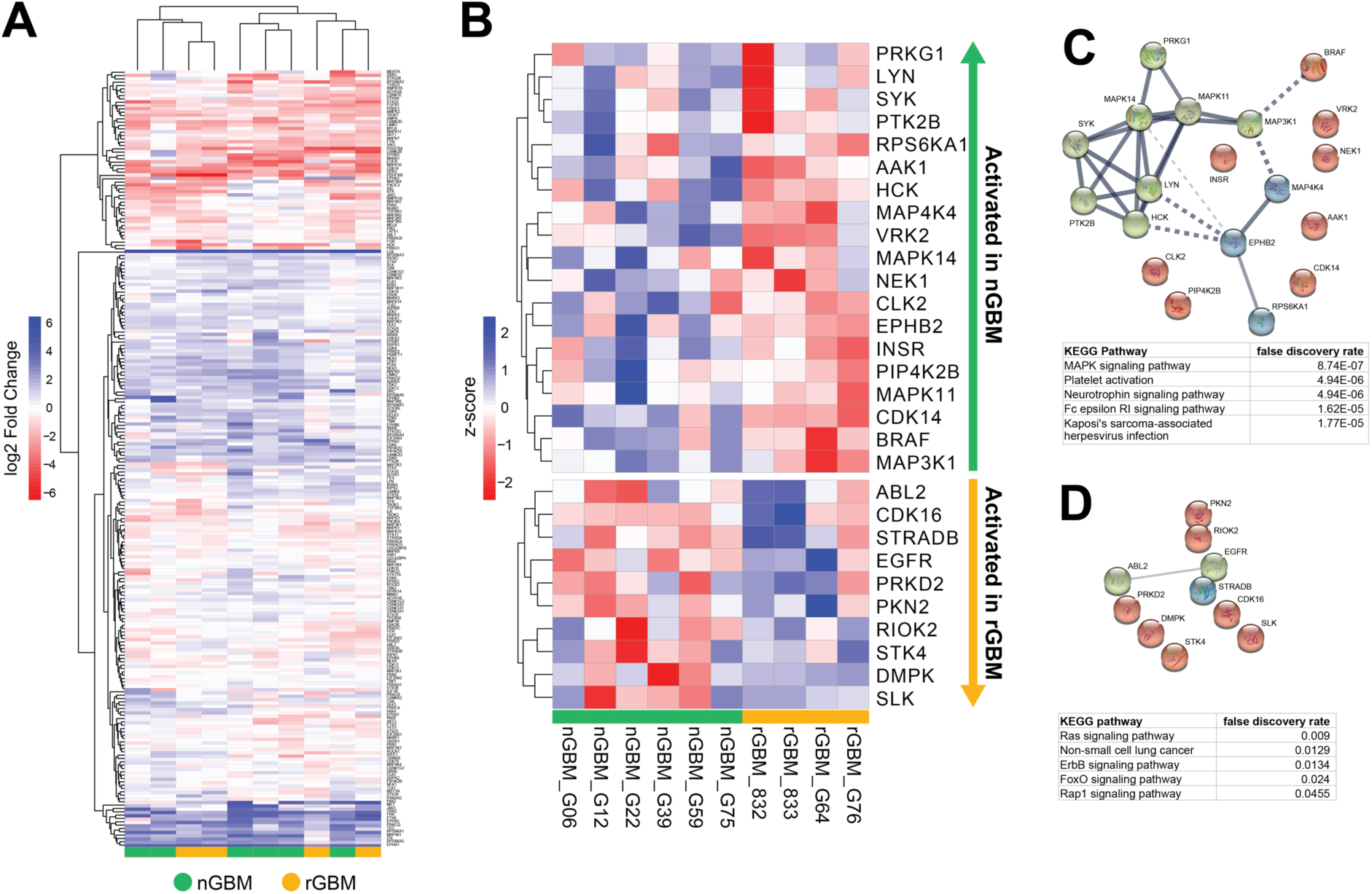
Kinome proteomic profiling of newly diagnosed and recurrent GBM reveals differences in kinase networks. **A**. log2FoldChange values for each kinase across six newly diagnosed and four recurrent GBM PDX tumors were calculated and visualized on a heatmap. **B**. Statistically altered kinases in nGBM or rGBM were identified (ANOVA, BH < 0.05). Heat map of row z-scores for each kinase. **C-D**. Protein networks and KEGG pathway analysis for kinases was performed for newly diagnosed (C) and recurrent (D) GBM.

In nGBM, statistical analysis reveals a tightly connected network of MAPK signaling that may drive tumor cell proliferation and migration upon tumor initiation (Figure 1C). The majority of kinases upregulated in nGBM are members of the tyrosine kinase (TK) or CMGC group, including several members of the MAPK signaling pathway. Two members of the Src kinase TK family, LYN and HCK, are activated, which have both been shown to have roles in suppressing apoptosis in GBM tumors.^29,30^. KEGG pathway enrichment analysis shows several key differences in tumorigenesis pathways. Significant neurotrophin signaling, driven by several members of the MAPK pathway and upstream regulators, may be responsible for the dramatic level of invasion seen in GBM tumors ^31^. Interestingly, two pathways related to inflammation were highly indicated in nGBM: Fc epsilon RI signaling and platelet activation. Fc epsilon RI is a surface protein expressed on platelets and has been previously implicated in cancer metastasis ^32,33^. Kinases enriched in inflammation pathways included LYN, SYK, and two p38 MAPK proteins. Importantly, LYN has been previously shown to activate SYK, leading to activation of platelets ^34^.

In rGBM, fewer tyrosine kinases were upregulated, however there was a shift towards serine/threonine (STE) kinases, including STK4, STRAD8, and SLK, which are members of the STE20 family. STRAD8 has been previously shown to drive GBM progression and elevated cytoplasmic levels is associated with decreased survival ^35^. Kinase clustering and KEGG pathway analysis reveals more active Ras and Erbb signaling in rGBM tumors, driven by the tyrosine kinases EGFR and ABL2, as well as STK4. This suggests a dependence on EGFR and downstream Ras signaling to promote tumor cell proliferation ^36,37^ (Figure 1D). Two pathways were enriched in both nGBM and rGBM: Rap1 signaling and FoxO signaling. Activation of Rap1 signaling, driven by EGFR and PRKD2 suggests a higher dependency on Rap1 for migration and invasion in rGBM ^38,39^. FoxO signaling, which is activated downstream of Erbb2, has been shown to promote GBM stem-cell like differentiation and is associated with tumor progression ^40,41^.

### RNA sequencing predictions of activated kinases

Differentially upregulated genes for GBM PDX lines were identified and significantly enriched gene ontology (GO) biological processes were identified. A two-way ANOVA (BH < 0.05) of GO terms showed that no significantly different ontologies were found between nGBM and rGBM PDX lines (Supplemental Table 1). Consistent with previous studies, transcriptional profiles alone using bulk RNA-sequencing does not provide insight into disease progression upon recurrence in GBM^42^

Active kinase predictions were made based on significantly upregulated mRNA for each GBM PDX line using the X2K platform. Using a two-way ANOVA (BH <0.05), X2K predicted kinases were identified, however all kinases that differed statistically were more active in newly diagnosed GBM (Figure 2A). STRING network of X2K predicted kinases shows a tightly clustered group of MAPK signaling proteins, as seen in kinome MIB-MS proteomic analysis (Figure 2B). Additionally, a network of CDK signaling may indicate dysregulation of cell cycle checkpoints, as previously reported in GBM ^37^. KEGG pathway enrichment of X2K predicted kinases was compared to KEGG pathways found using MIB-MS technology. While all pathways found to be upregulated in the MIB-MS analysis of nGBM were recapitulated in the nGBM X2K predicted kinases analysis, nearly half of pathways activated in the X2K dataset were not found using the MIB-MS technology (Figure 3A-B). Furthermore, X2K nGBM kinases showed an enrichment of Erbb signaling and Ras signaling, as seen in pathway analysis for the rGBM kinome profiles (Figure 3C), indicating that the X2K kinase subset may not be specific for newly diagnosed GBM.

**Figure 2.**
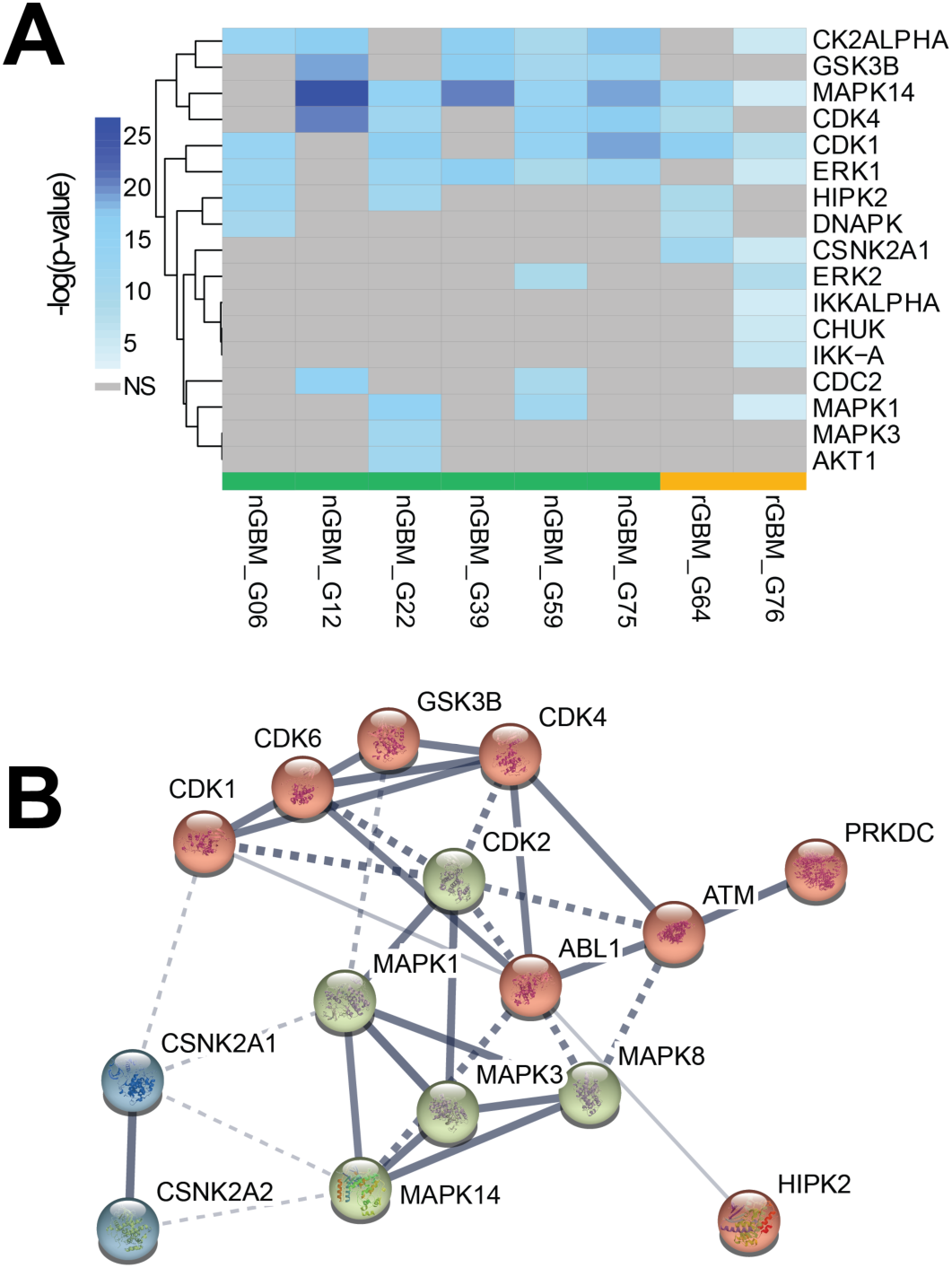
X2K predicted kinases activated in nGBM show MAPK signaling enrichment. Genes significantly upregulated in GBM PDX tumors were used as input for the X2K platform to identify kinases predicted to be active. **A**. Significantly activated X2K kinases in nGBM (ANOVA BH < 0.05) were identified and plotted on a heatmap. **B**. STRING network illustrates protein-protein interactions of X2K activated kinases in nGBM.

**Figure 3.**
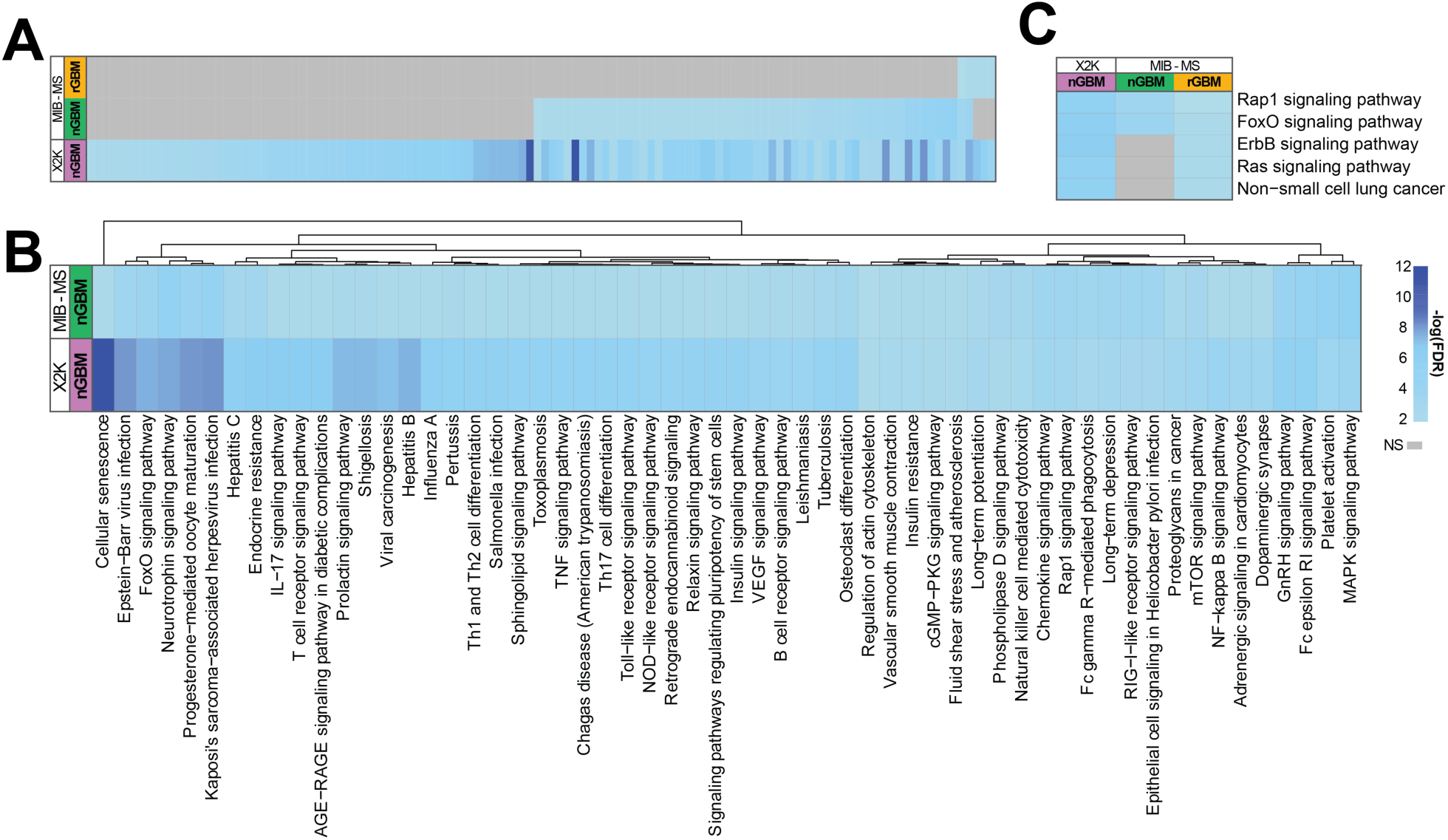
X2K predictions for active kinases show different signaling pathways from kinases identified using MIB-MS. **A**. KEGG pathways enriched with altered kinases in the MIB-MS nGBM and rGBM, as well as X2K nGBM were identified and –log(FDR) values were plotted on a heat map (FDR > 0.05). **B**. Pathways enriched for nGBM MIB-MS kinases are plotted alongside pathways in nGBM X2K kinases. **C**. Pathways enriched in MIB-MS rGBM are plotted alongside nGBM MIB-MS and nGBM X2K pathways.

## DISCUSSION

Here we report several kinome reprogramming mechanisms seen in newly diagnosed and recurrent glioblastoma. Kinases differentially upregulated or downregulated in nGBM and rGBM were identified using MIB-MS, providing a unique view of the tumor kinome that arises upon GBM tumor recurrence. We were able to see changes in kinases driving invasion and cell differentiation, as well as alterations in the tumor microenvironment. Analysis of RNA-sequencing for the same panel of tumors was unable to capture the same level of complexity in the kinome reprogramming.

Glioblastoma is marked by a high degree of inter- and intra-tumor heterogeneity. Despite a significant difference in survival between newly diagnosed and recurrent GBM, transcriptional profiling of tumors separated by disease status yields minimal patterns in gene ontology or pathway analysis (Supplemental Table 1). Upon tumor recurrence, exome sequencing has shown that many somatic mutations are not conserved and transcriptional subtype is often not maintained, suggesting a high degree of evolution from the initial tumor ^19,43^. Additionally, drug sensitivity between patients varies widely ^16^.

Here we report a distinct kinase protein signature for newly diagnosed and recurrent glioblastoma in a panel of six nGBM and four rGBM PDX tumors. Differences in invasion-related pathways were identified, with a shift away from neurotrophin activation in nGBM to Rap1 activation in rGBM. Our results also suggest that platelet involvement in nGBM, which has been implicated in tumor growth, may be mediated by kinases^44^. Furthermore, platelet activation has been implicated in GBM tumor growth ^44^. Previous studies have found that levels of sphingosine-1-phosphate (S1P), a lipid signaling molecule secreted by platelets that has been implicated as a modulator of inflammation and tumorigenesis, is significantly elevated in GBM tumors ^45^. S1P drives the transformation of macrophages into tumor associated macrophages (TAMs), which play a role in driving tumor cell invasion by suppressing the immune system in the tumor microenvironment ^45^. Platelet activation has also been shown to drive tumor invasion in other cancers, and kinases mediated platelet activation has been reported ^46-48^. Additionally, pathways involved in cell differentiation in recurrent GBM were identified. Erbb2 signaling may lead to downstream FoxO activation, which promotes glioblastoma stem-cell like (GSC) tumor cell differentiation upon tumor recurrence. GSCs have been heavily implicated as drivers of intra-tumor heterogeneity and tumor recurrence ^49^. Targeting kinases involved in GSC tumorigenesis provides a potential therapeutic strategy for future studies.

At a transcriptional level, significantly enriched GO biological processes for differentially expressed genes did not show any differences between nGBM and rGBM. Additionally, active kinase predictions from the X2K platform showed little overlap with the kinase proteome signatures. Using a KEGG analysis of kinases predicted to be activated in nGBM or rGBM, we were unable to identify relevant pathways that differed between disease states. We also identified statistically different kinases between nGBM and rGBM, however only kinases statistically significantly activated in nGBM were identified. These kinases formed a tight network of MAPK signaling proteins, as seen in the nGBM kinase proteome profile. General MAPK signaling pathways were evident in both transcriptional and proteomic kinase analyses.

Prior studies have shown the heterogeneity of the kinome in GBM, and that tumors can be stratified based on kinome profile. Novel resistance mechanisms exist based on kinome profile, however as we have shown here, some kinase pathways may be conserved across multiple GBM tumor types upon recurrence ^22^. As described previously, transcriptional data does not provide enough information for molecular studies in GBM, while proteomic studies are able to provide more relevant pathways to inform future therapies ^50,51^.

MIB-MS kinome signatures can provide insight into RTK mechanisms underlying tumor recurrence in GBM. As contrasted with transcriptional profiling, measuring levels of kinases using MIB-MS for nGBM and rGBM provides a more robust kinome signature. Kinase proteomic profiling was able to elicit a strong signature for both newly diagnosed and recurrent GBM that is not evident at a transcriptional level. However, transcriptional levels were unable to provide significant insight into the difference between kinases involved in newly diagnosed and recurrent GBM. Collectively, our studies suggest that kinase proteomic profiles may provide a means of identifying therapeutic targets in GBM by tumor type, as well as provide a potential mechanism for tumor recurrence.

## Abbreviations

GBM: Glioblastoma
MIB-MS: Multiplexed Inhibitor Beads and Mass Spectrometry
TMZ: Temozolomide

## ACKNOWLEDGMENTS

We thank all members of the Center for Therapeutic Innovation and the Lemmon-Bixby laboratories for helpful discussions. We acknowledge support from the Sylvester Comprehensive Cancer Center, and R01 CA211670 (J.S.D.), NIH T32 CA009035 (A.M.K). N.G.A. is a consultant for Epigenetix, Inc.

